# Rewiring Dynamics of Functional Connectome in Motor Cortex during Motor Skill Learning

**DOI:** 10.1101/2022.07.12.499746

**Authors:** Saber Meamardoost, EunJung Hwang, Mahasweta Bhattacharya, Chi Ren, Linbing Wang, Claudia Mewes, Ying Zhang, Takaki Komiyama, Rudiyanto Gunawan

## Abstract

The brain’s connectome continually rewires throughout the life of an organism. In this study, we sought to elucidate the operational principles of such rewiring by analyzing the functional connectomes in mouse primary motor cortex (M1) during a 14-session (day) lever-press task learning in response to an auditory cue. Specifically, we employed Calcium imaging recordings of L2/3 and L5 of M1 in awake mice to reconstruct and analyze functional connectomes across learning sessions. Our results show that functional connectomes in L2/3 and L5 follow a similar learning-induced rewiring trajectory. More specifically, the connectomes rewire in a biphasic manner, where functional connectivity increases over the first few learning sessions, and then, it is gradually pruned to return to a homeostatic level of network density. We demonstrated that the increase of network connectivity in L2/3 connectomes, but not in L5, generates neuronal co-firing activity that correlates with higher motor performance (shorter cue-to-reward time), while motor performance remains relatively stable throughout the pruning phase. The results show a biphasic rewiring principle that involves the maximization of reward / performance and maintenance of network density. Finally, we demonstrated that the connectome rewiring in L2/3 is clustered around a core set of movement-associated neurons that form a highly interconnected hub in the connectomes, and that the activity of these core neurons stably encodes movement throughout learning.

**Significance Statement:** Connectomes in the motor cortex rewire during motor skill learning, but the operational principle behind this rewiring is yet to be determined. Here, we characterized the rewiring dynamics of functional connectomes in L2/3 and L5 of M1 in mice engaging in a lever-press learning, using two-photon fluorescence microscopy data. We identified a universal biphasic rewiring trajectory across animals and layers in the motor cortex that reflects two objectives: an exploratory phase that increases functional connectivity and optimizes motor performance, and a pruning phase that brings connectivity back to a homeostatic level while maintaining motor performance. We found further that connectome rewiring during motor skill learning concentrates around a core set of highly interconnected neurons in L2/3 that reliably encode movements.

## Introduction

The synaptic wiring of neurons, also known as connectome, is the foundation of brain’s myriad functions. The brain’s connectome is not static; quite the opposite, it continues to rewire throughout an organism’s life (Citri and Malenka, 2008; Bassett et al., 2011; Caroni et al., 2012; Bennett et al., 2018; Papale and Hooks, 2018a; Kao et al., 2020). Synaptic plasticity is critical in enabling organisms to learn new skills, store new memory, and adapt and respond appropriately to novel environment (Bassett et al., 2011; Caroni et al., 2012; Papale and Hooks, 2018a; Kao et al., 2020). Understanding the principles of how the brain’s connectome rewires may facilitate, not only the formulation of therapies for neurological disorders, but also the development of novel paradigms of data storage and computing architecture (Bathe et al., 2021).

Many studies have shown substantial synapse formation and connectome rewiring in primary motor cortex (M1), particularly in layers 2/3 and 5 (Greenough et al., 1985; Withers and Greenough, 1989; Wang et al., 2011). Synaptic reorganization may occur through formation and degradation of synaptic connectivity between pre- and post-synaptic neurons, and the associated appearance and disappearance of dendritic spines (Fu et al., 2012; Peters et al., 2014; Huang et al., 2020; Albarran et al., 2021). Spine formation has been observed across a variety of motor learning tasks (Papale and Hooks, 2018b) such as reaching and grasping (Xu et al., 2009), lever-pressing (Peters et al., 2014), and running on an accelerated motorized rod (Yang et al., 2009), across M1 layers, including L2/3 (Peters et al., 2014) and L5 (Xu et al., 2009; Yang et al., 2009). fMRI recording of M1 in adult humans previously showed two separate phases during a finger tapping task, responsible for within- and across-session learning (Kami et al., 1995). During the first training session and after a few trials, a small region in M1 is activated and shows a ‘habituation-like’ activity for fast learning. However, during the later sessions a larger area in M1 is activated for long-term reorganization of M1 for motor learning. While progress has been made in understanding how the circuitry of brain regions reconfigures during learning (*e*.*g*., using functional MRI (fMRI) data (Bassett et al., 2011; Kao et al., 2020)), the principles behind functional rewiring among individual neurons are much less understood. Recent advances in neural activity recordings, such as two-photon calcium imaging (Stosiek et al., 2003), have produced large-scale firing data of single neurons in awake animals while learning various tasks (Peters et al., 2014, 2017), opening the avenue to study brain’s connectome at neuronal population level.

This work focuses on the functional connectomes in L2/3 and L5 of M1. The aim of our study is to elucidate how functional connectomes in these regions are rewired during a motor-task learning and what operational objective(s) drives the connectome rewiring. To this end, we reconstructed functional connectomes from two-photon Calcium (Ca^2+^) recordings and characterized the connectome rewiring dynamics during learning of a lever pressing task in response to an auditory cue (Peters et al., 2014, 2017). We analyzed how functional connectomes in both layers of M1 rewire during learning and how this rewiring is associated with motor performance. We then studied the rewiring of neurons that are most impacted by learning and identified groups of neurons that have different learning-induced rewiring patterns. Our results show that the functional connectomes in both layers continually rewire throughout the learning period following a common trajectory where connectomes transiently increase their functional connections before returning to a homeostatic level of connectivity. Further, we observed the existence of an interconnected hub of movement-associated neurons in the connectomes that are heavily rewired and stably encode movement during learning.

## Methods

### Animals

Two groups of mice were used in this work from two separate studies previously published where neuronal activity in L2/3 (Peters et al., 2014) and L5 (Peters et al., 2017) was recorded during learning a lever-press task. These animals are referred to as L2/3 (*n =* 7) and L5 (*n =* 8) mice corresponding to the cortical layer of neurons from each mouse. Note that one animal (Mouse L2/3-6) was removed from the analyses due to multiple missing sessions of training.

### Experiments: lever-press task

Details on experimental procedure and data collection available in the original publication (Peters et al., 2014, 2017). Briefly, genetically encoded Calcium indicator GCaMP5G was virally expressed in the motor cortex of mice (C57BL/6). In the experiments, water deprived head-fixed mice performed a lever-press task learning. In the task, the mice learned to press a lever beyond a set threshold (∼ 3 mm) using their left forelimb within 10-30 seconds after an auditory tone was presented in order to receive a water droplet as reward (see **Fig. 2a**). Calcium fluorescence images of L2/3 and L5 neuronal activity were recorded at ∼28 Hz using two-photon microscopy, from the same field of view (472 × 508 *μm* area) over the 14 learning sessions (1 session/day; 20-30 minutes/session). Meanwhile, lever displacement trace was recorded at 10 kHz. Regions of interest (ROIs) in the Calcium images were manually drawn to demarcate the somas or dendritic shafts of individual neurons and then, aligned across sessions. Pixels within each ROI were averaged, and background fluorescence fluctuations were subtracted from the average to create a Calcium fluorescence time series (dF/F) for each neuron.

### Functional connectome inference

Functional connectomes were inferred from two-photon (2p) Ca^2+^ fluorescence imaging data using FARCI, a recently developed connectome inference pipeline (Meamardoost et al., 2021). Briefly, FARCI uses Online Active Set method to Infer Spikes (OASIS) algorithm (Friedrich et al., 2017) from the Suite2p package (Pachitariu et al., 2018), to deconvolve neuronal spiking activity from 2p Ca^2+^ imaging data. Subsequently, the deconvolved spikes are thresholded to keep only significant Ca^2+^ spikes (larger than *μ* + 2*σ*) and then, smoothened using a moving window weighted average (5 frames) (Meamardoost et al., 2021). Finally, a partial correlation matrix is generated to provide the adjacency matrix of the functional connectome. In order to remove spurious connections, we applied a thresholding strategy, keeping only the partial correlation coefficients that are larger than *μ* + 2*σ* in each session. Each element in this matrix represents the functional connectivity between two corresponding neurons, and each pair of neurons forms an edge in the connectome.

### Functional connectome rewiring

To study the dynamics of connectome rewiring over multiple sessions of motor skill learning, the functional connectome inference as described above was applied to the neuronal activity data collected from different sessions separately. This step generated an (*n* × *n*) adjacency matrix, where *n* is the number of neurons in the population, for each learning session. The upper triangular part of this matrix was extracted and stored as a single-row adjacency vector containing 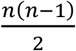 elements. The adjacency vectors from different sessions were then stacked to give a matrix with 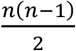 columns and *s* rows, where *s* is the number of sessions. Finally, Principal Component Analysis (PCA) was applied to this matrix—each row vector from a session represents an observation, while each column vector (partial correlation between two neurons) is a feature. The PC scores were used to visualize the connectome rewiring dynamics during motor skill learning.

### Trial-based functional connectome activity

To evaluate the activity of functional connectome in a rewarded trial, we first extracted smoothened Ca^2+^ spikes from movement-related frames in the respective session, as illustrated in **Fig. 1a**. We applied FARCI to generate movement-related functional connectome for the learning session. The adjacency matrix of this connectome was then binarized—partial coefficients that are larger than *μ* + 2*σ* are set to 1, and otherwise 0. Separately, we extracted smoothened Ca^2+^ spikes for each rewarded trial and evaluated pairwise Pearson’s correlations for all neuron pairs to produce a correlation matrix (see **Fig. 1b**). Finally, we computed the Hadamard product of the Pearson correlation matrix of a trial and the binarized adjacency matrix of movement-related functional connectome of the session, to give trial-based activity of the connectome (see **Fig. 1c**). Following the visualization of functional connectome rewiring above, we extracted the upper triangular part of the resulting connectome activity matrices and applied PCA to obtain a reduced dimensional visualization.

**Figure 1.**
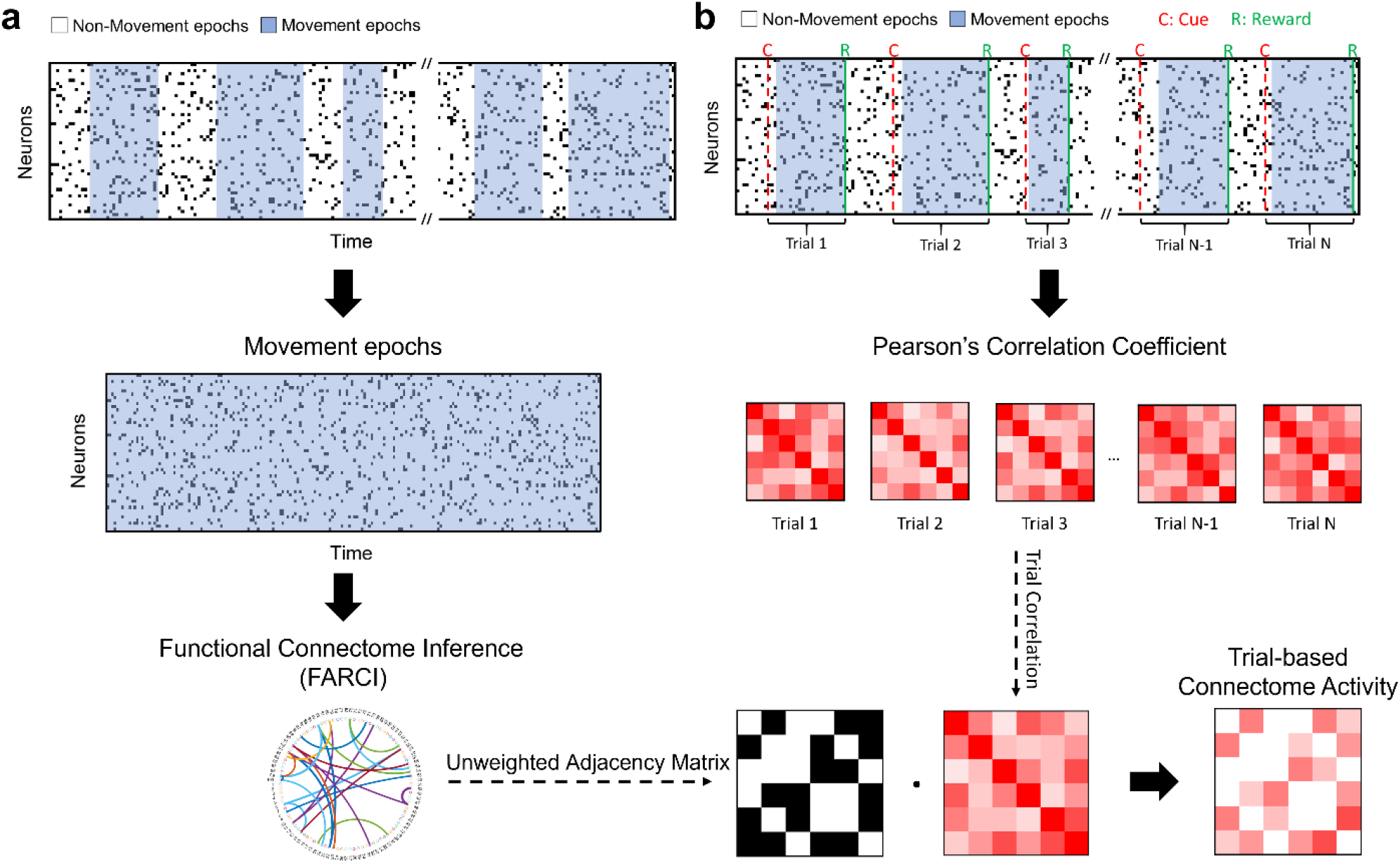
Data processing for the inference of functional connectome and trial connectome activity. **a**. Functional connectome inference pipeline. For each session, movement epochs are extracted from the neuronal activity data and FARCI is applied to produce the movement-related functional connectome. **b-c**. Trial-based connectome activity. Pairwise Pearson’s correlations are computed using the activities of neuron pairs for a rewarded trial (cue-to-reward), which are then filtered using a binary adjacency matrix obtained from functional connectome, to give the connectome activity of the trial.

### Time-series PCA

Time-series PCA was applied to project neuronal population activity during rewarded trials to a lower dimensional space (Sauerbrei et al., 2020). To do so, for each rewarded trial, we identified the movement epoch immediately before the reward, and set the first frame of this epoch as the movement onset time (see Fig. 1b). We extracted smoothened Ca^2+^ spikes from a 98-frame time window that was defined such that the movement onset is the 15-th frame in this window. We applied the following time-series PCA for each session separately.

First, we converted the smoothened Ca^2+^ spikes of each neuron to z-scores by using their trial-based average and variance. These z-scores were then averaged across the trials in a session, to which the standard PCA was applied—each time point was an observation and the average z-score of a neuron was a feature. The resulting principal components were then used to visualize the time-series neuronal activity as 3D trajectory in PC1-3 space.

### Network connectivity analysis

We employed four different metrics: mean degree, network density, transitivity, and clustering coefficient, to characterize the interconnectedness of the functional connectomes. Mean degree is the average number of connections that a neuron has. Meanwhile, network density gives the ratio of the number of existing connections and the total possible number of connections. Transitivity is computed as the ratio between the number of existing closed triplets in the connectome graph and the total possible number of triplets in a neuronal population. Finally, clustering coefficient is defined for each node (neuron) *i* as the ratio between the number of triangles connected to neuron *i* and the number of triplets centered on *i*. For a functional connectome, we evaluated the mean (average) clustering coefficient across all neurons. Both transitivity and clustering coefficients give a measure for the tendency of neurons to form clusters in the connectome. For all of the above metrics, a higher value indicates a more interconnected network.

### Classification and analysis of movement-related neurons

We followed the procedure outlined in the original publication to classify neurons as ‘movement-related’ (Peters et al., 2014). The classification was performed based on the level of neuronal activity during movement epochs. Briefly, binarized lever traces were used to label the Ca fluorescence imaging frames into movement and quiescent epochs. The mean activity of each neuron was computed over the movement epochs in a session. Subsequently, the movement and quiescent epochs were shuffled, thereby randomizing the relative position of these epochs with each other. The mean activity of neurons for the shuffled epochs was also evaluated, and this shuffling was repeated for 10,000 times. A neuron was classified as movement related if its mean activity during the movement epochs was higher than the 0.5th percentile of the mean activities computed for the randomly shuffled epochs.

To evaluate the fraction of movement related neurons in the edges that account for the largest connectome changes over learning, we first sorted edges based on their loadings to PC1 in the above PCA of functional connectome rewiring in descending order. Subsequently, we identified two groups: the highest 100 edges (top positive PC1 loadings) and the lowest 100 edges (top negative PC1 loading). For each group, we compiled neurons that are incident to these edges—two for every edge--resulting in a set of 200 non-unique neurons for each group. For each neuron in these groups, we calculated the number of times that it was classified as movement-related across different periods of learning: sessions 1-2, 3-5, 6-9, and 10-14. The fraction of movement-related neurons was computed by dividing the total number of times that the top PC1 neurons are classified as movement-related with the total number of neurons multiplied by the number of sessions in the learning period (i.e., 200 x *m*, where *m* is the number of sessions in a learning period. For example, for sessions 6-9, *m* is 4). To establish the significance level, we repeated the above procedure using 100 edges that are randomly sampled from the union of edges in the connectomes across all sessions, for a total of 1000 times. To establish statistical significance, we evaluated the *k*-th upper percentile associated with the fraction of movement-related neurons for each learning period with respect to the distribution of the fraction from the random sampling of edges.

### Evaluation of session-to-session changes in functional connectivity

To assess the degree to which neurons change their connectivity during motor learning, we first computed functional connectome in each session and evaluated the number of connections that change between two consecutive sessions for each neuron. Next, for a given group of neurons, we computed the average session-to-session change in connectivity as the ratio of total number of altered connections across all the neurons to *n* × (*s* − 1) where *n* and *s* are the number of neurons and sessions, respectively.

### Identification of Core, NP, EP, and O neurons

We classified neurons based on their connectivity changes into four types: Core, Naïve Phase (NP), Expert Phase (EP), and Other (O). The classification used the PCA of the functional connectomes described above. For each mouse, we determined the set of edges in the functional connectome that are associated with highly positive and negative PC1 loadings using a threshold determined by the knee detection algorithm (Satopaa et al., 2011). Then, we identified two sets of neurons, one set that is incident to connectivity from the top positive PC1 loading and another to the top negative PC1 loadings. Neurons belonging to both sets are labelled as Core neurons. Those that are associated with only the top positive PC1 loadings are labeled as EP, while those associated with only the top negative PC1 loadings as NP. Neurons that are not in the three sets above are called Other (O) neurons.

### Correlations with expert (learned) activity and movement pattern

To examine the similarity of neural population activity in an individual trial to the learned expert activity or the similarity of movement to the learned movement pattern, we performed correlation analysis. Neuronal spiking activity data were preprocessed as above. The expert activity and movement patterns refer to the average activity and movement in 84-frame windows of rewarded trials over sessions 10-14. The first frame in this 84-frame window corresponds to the movement onset.

### Linear decoders of neuronal activity to lever position

To map neuronal activity to movement for each group of neurons, we used pre-processed neuronal spiking activity data as above. Rewarded trials in each session were then split by random into train and test sets at 80% and 20% ratio, respectively. For a given session, a decoder was trained using linear regression (ordinary least square) to predict lever position (*i*.*e*., lever voltages) from the spike data of a specific set of neurons (Core, EP, NP, and O neurons). The performance of the trained linear decoder was then assessed by computing the correlations between the predicted and actual lever positions (measured in voltages) using data from the test set. In addition, an expert decoder was trained using the combined data from sessions 10-14, and its performance was tested on the data from sessions 1-3.

## Results

### Motor skill learning reconfigures functional connectome in the motor cortex

Lever-press task learning experiments were previously carried out where water-deprived head-fixed mice (*n =* 7) learned to press a lever in response to an auditory cue in order to receive a water droplet reward. Neuronal activity from L2/3 or L5 of mouse primary motor cortex (M1) were recorded using 2p Ca^2+^ imaging (see illustration in **Fig. 2a**) (Peters et al., 2014, 2017). The average times between the cue and the corresponding reward decreased with learning (see **Fig. 2b**), indicating that these mice were able to acquire reward efficiently.

**Fig. 2.**
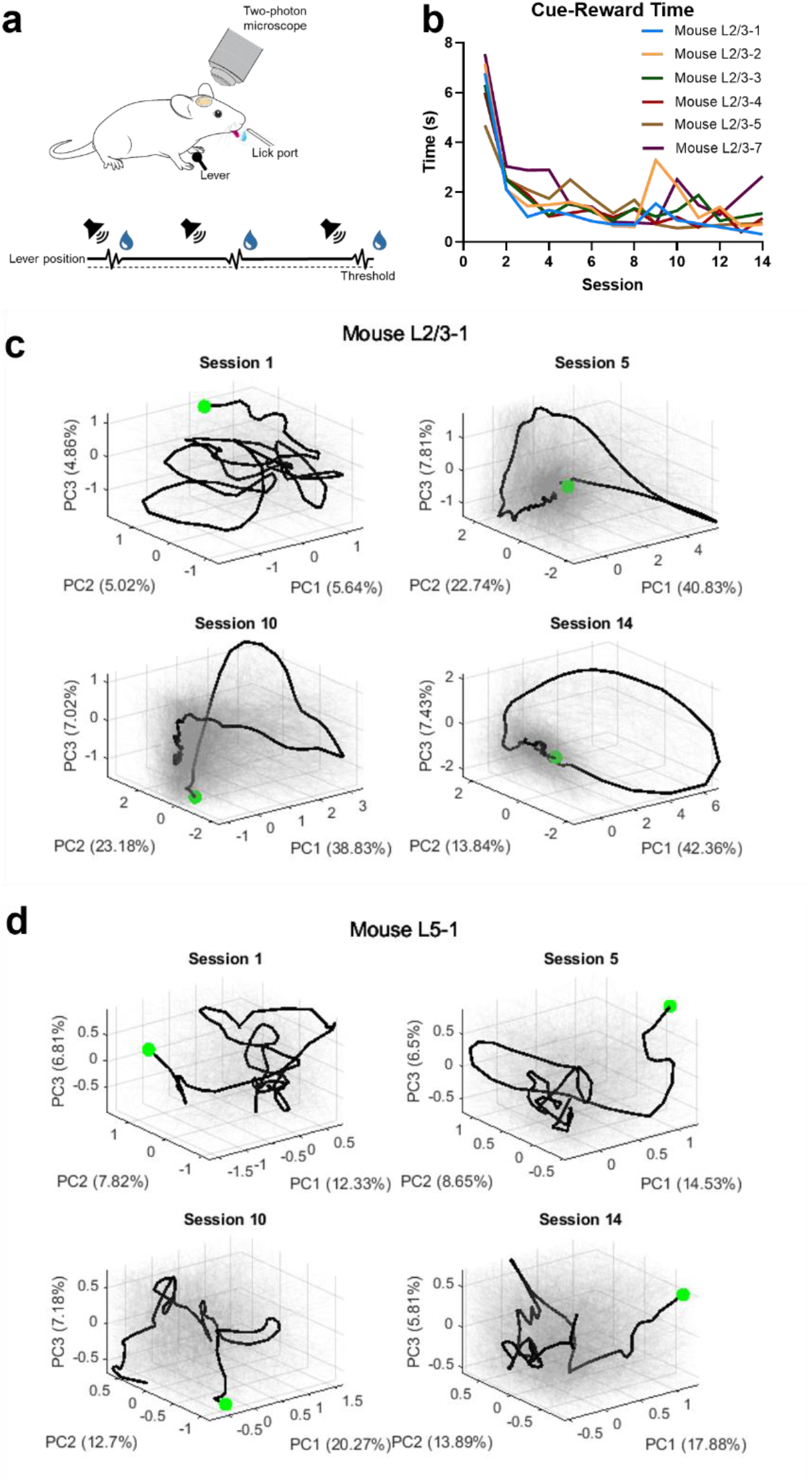
Neuronal population activity of L2/3 and L5 regions of primary motor cortex M1 in lever-press task learning. **a**. Lever-press task schematic. **b**. Mean duration between cue and reward, averaged across all rewarded trials from each session. **c, d**. Scores of the first three principal components (PCs) of neuronal population activity across sessions in L2/3 (Mouse L2/3-1) and L5 (Mouse L5-1) regions, respectively. The neuronal population activities were extracted from a 98-frame window of every rewarded trial. The window was anchored by the movement onset time, which was set to be the 15^th^ frame in this window. The green dot marks the starting time point.

Lever-press task learning reshaped the neuronal activity in M1 L2/3 generating reproducible neuronal activity pattern and more consistent relationship between neuronal activity and movements (Peters et al., 2014). Time-series PCA of L2/3 neuronal activity in **Fig. 2c** confirms the emergence of temporal activity pattern with learning with increased synchronization of activity at or around the movement onset. In contrast, the activity of L5 corticospinal neurons did not become more consistent with learning. Rather, learning led to more dissimilar neuronal activity for dissimilar movements (Peters et al., 2017). In congruence with these observations, time-series PCA of L5 neuronal activity as shown in **Fig. 2d** shows a lack of trial-to-trial coherence that emerges with learning. Thus, lever-press task learning induces distinct reorganizations of neuronal population activities across different layers of M1.

To study functional connectome rewiring dynamics associated with lever-press task learning, we used partial correlations of neuronal spikes, computed using a recent connectome inference method FARCI, to establish functional connectivity among excitatory neurons in L2/3 and in L5. **Fig. 3a** displays L2/3 functional connectome rewiring, projected to the first two PC axes (see Methods), portraying homogeneous common trajectories for different animals. The rewiring trajectories show that the strongest contributor to the functional connectome variation across sessions (PC1) is associated with learning. **Fig. 3b-c** illustrate the dynamics of the average partial correlations associated with the 100 most positive and most negative loadings from the first and second PCs (*i*.*e*., PC1 and PC2). The dynamics indicates the continuous gain or loss of functional connections along the PC1 axis, and transitory gain or loss of connections along the PC2 axis. Finally, **Fig. 3d** illustrates the connectivity changes in the functional connectomes (mean degree, network density, transitivity, mean clustering coefficients) over the animals in the study, showing a sharp increase in the functional interconnectedness among neurons during the first 3-4 sessions, followed by a more gradual decrease in connectivity during the remaining sessions. The results above suggest a common rewiring dynamics of L2/3 functional connectomes among the mice subjected to lever-press task learning, where the functional connectome quickly increases its degree of connectivity (increased co-firing among neurons) during the initial stages of learning, and is then pruned to bring network interconnectedness toward the same level as that before learning.

**Figure 3.**
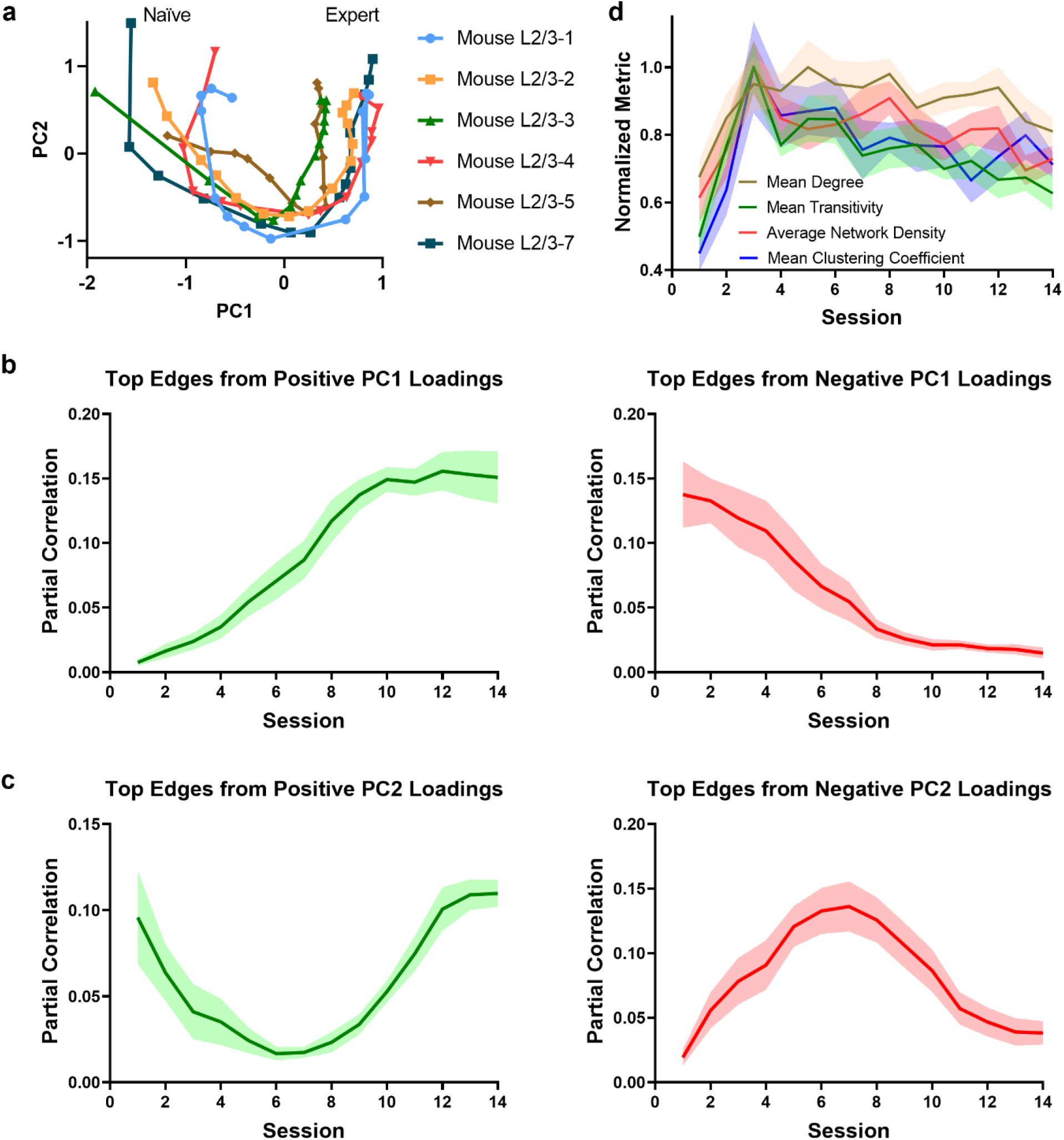
Functional connectome rewiring in L2/3 during lever-press task learning. The shaded areas indicate s.e.m., computed over the mice in the study. **a**. Connectome rewiring trajectories. Functional connectomes are projected onto PC1 and PC2 axes (see Methods). **b-c**. Average partial correlation coefficients of top 100 edges based on positive and negative magnitudes of PC1 and PC2 loadings. **d**. Network connectivity metrics of the functional connectomes. To aid comparison, normalized metrics are shown, using the highest value of each metric across sessions and animals as a scaling factor.

**Fig. 4a-c** gives the functional connectome rewiring dynamics for L5. Surprisingly, despite the differences between L2/3 and L5 in how lever-press task learning alters neuronal activities, similar rewiring trajectories are observed across all mice in these layers. The first PC axis is again associated the continuous gain or loss of functional connections in the connectome associated with learning, while the second PC axis captures the transitory gain or loss of functional connections. **Fig. 4d** gives the dynamic changes in network interconnectedness for the functional connectomes in L5, showing similar dynamics, albeit more subdued and delayed when compared with those in L2/3. The similarities in the functional connectome rewiring dynamics between L2/3 and L5 suggest the existence of a common operating principle.

**Figure 4.**
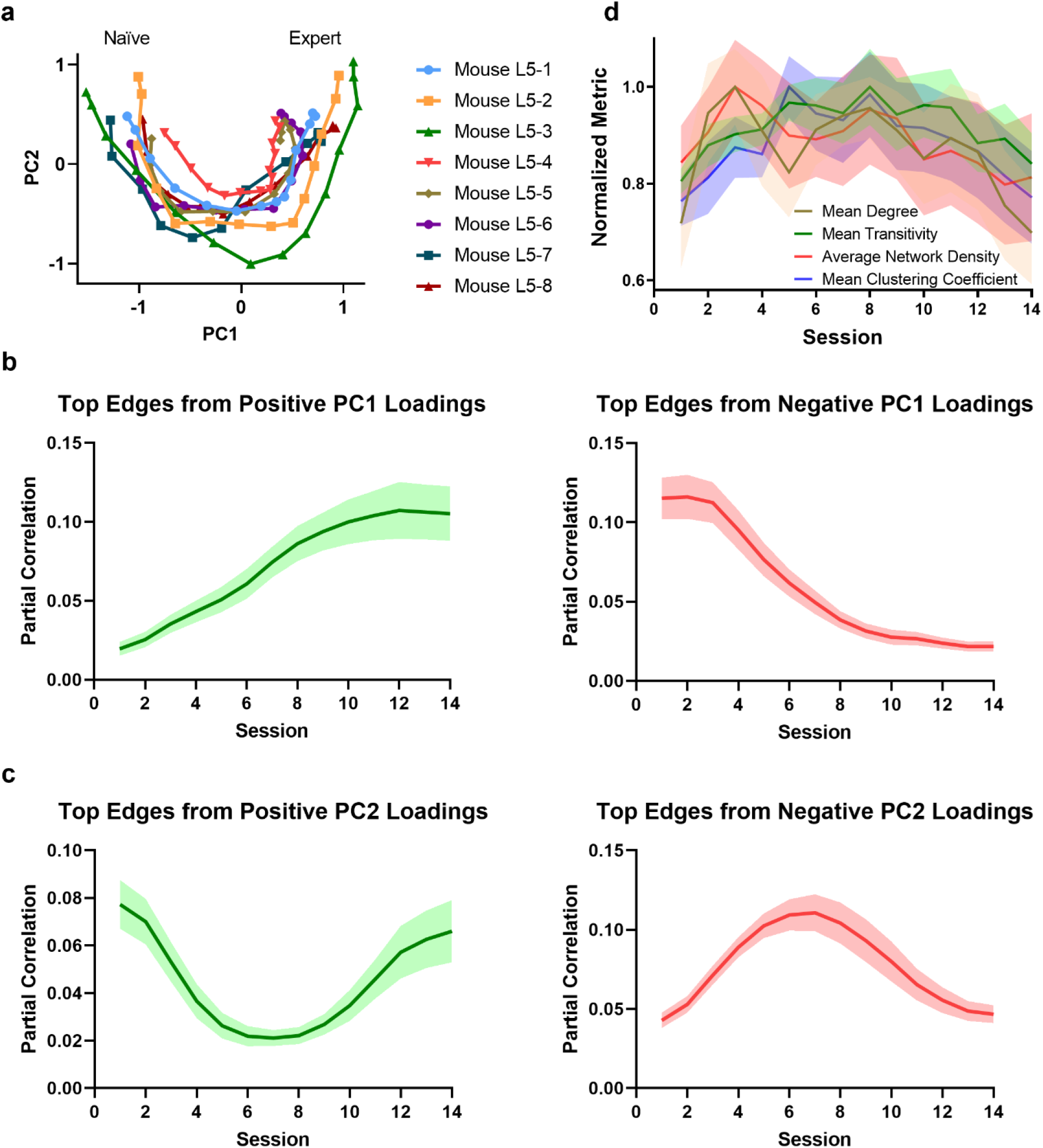
Functional connectome rewiring in L5 during lever-press task learning. The shaded areas indicate s.e.m., computed over the mice in the study. **a**. Connectome rewiring trajectories. Functional connectomes are projected onto PC1 and PC2 axes (see Methods). **b-c**. Average partial correlation coefficients of top 100 edges based on positive and negative magnitudes of PC1 and PC2 loadings. **d**. Network connectivity measures of the functional connectomes. To aid comparison, normalized metrics are shown, using the highest value of each metric across sessions and animals as a scaling factor.

### Connectome rewiring in motor cortex is a biphasic process

Next, we studied how functional connectome activity is related to motor task performance on a trial-by-trial basis. To this end, we evaluated functional connectome activities using gated pairwise Pearson correlations of neuronal Ca^2+^ spike activities in rewarded trials (see Methods), providing a measure for the co-firing activity of neurons. **Fig. 5** depict the L2/3 functional connectome activities from rewarded trials in individual mice, and their associated cue-to-reward times. Expectedly, the functional connectome activities across sessions follow a similar trajectory in the PC1-PC2 projection (*x-y* axes) to the rewiring dynamics shown above. We noted that the relationship between L2/3 functional connectome activity and task performance can be divided into two phases: in the first phase (up to session 3-4), alterations in the functional connectome activities are associated with a rapid improvement in motor performance leading to a sharp drop in the cue-to-reward time. In the subsequent phase, functional connectome activities continue to change while motor performance remains stable. Interestingly, as shown in **Fig. 3a** & **d**, the first phase is marked by a sharp increase in the connectivity of the functional connectomes, while the second phase corresponds to a period of gradual pruning of the connectome. **Fig. 6** shows the L5 functional connectome activity and the motor performance for every rewarded trial from individual mice. Consistent with the key findings of the original study (Peters et al., 2017), the activity of L5 functional connectomes has a weaker relationship with the motor performance in comparison with L2/3.

**Figure 5.**
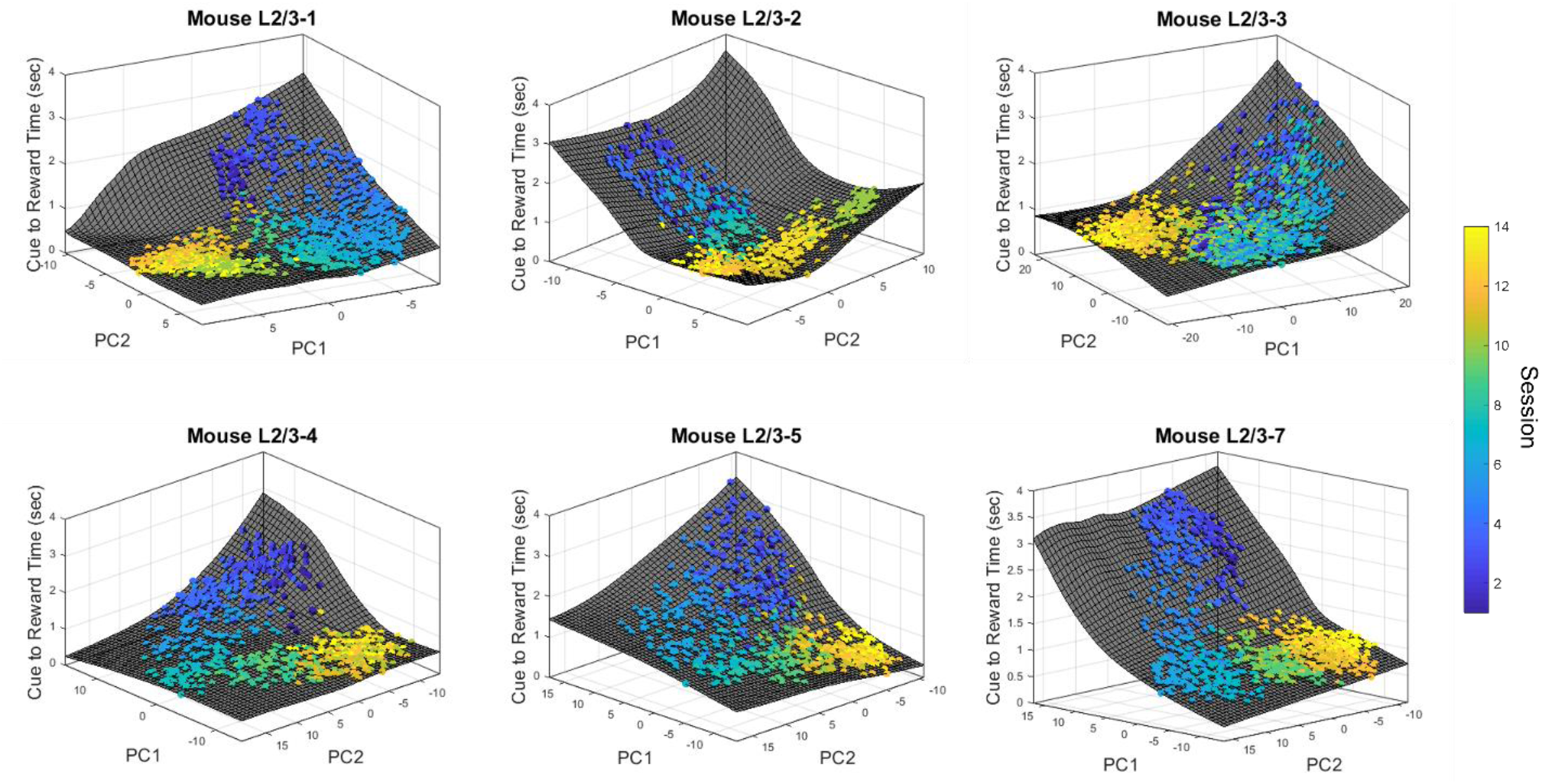
L2/3 connectome activity versus cue-to-reward time for rewarded trials. Each circle represents a rewarded trial and its color denotes the session number. The surface interpolation was done using MATLAB function `cftool`. The cue-to-reward time of each trial is projected onto the surface for visualization purposes.

**Figure 6.**
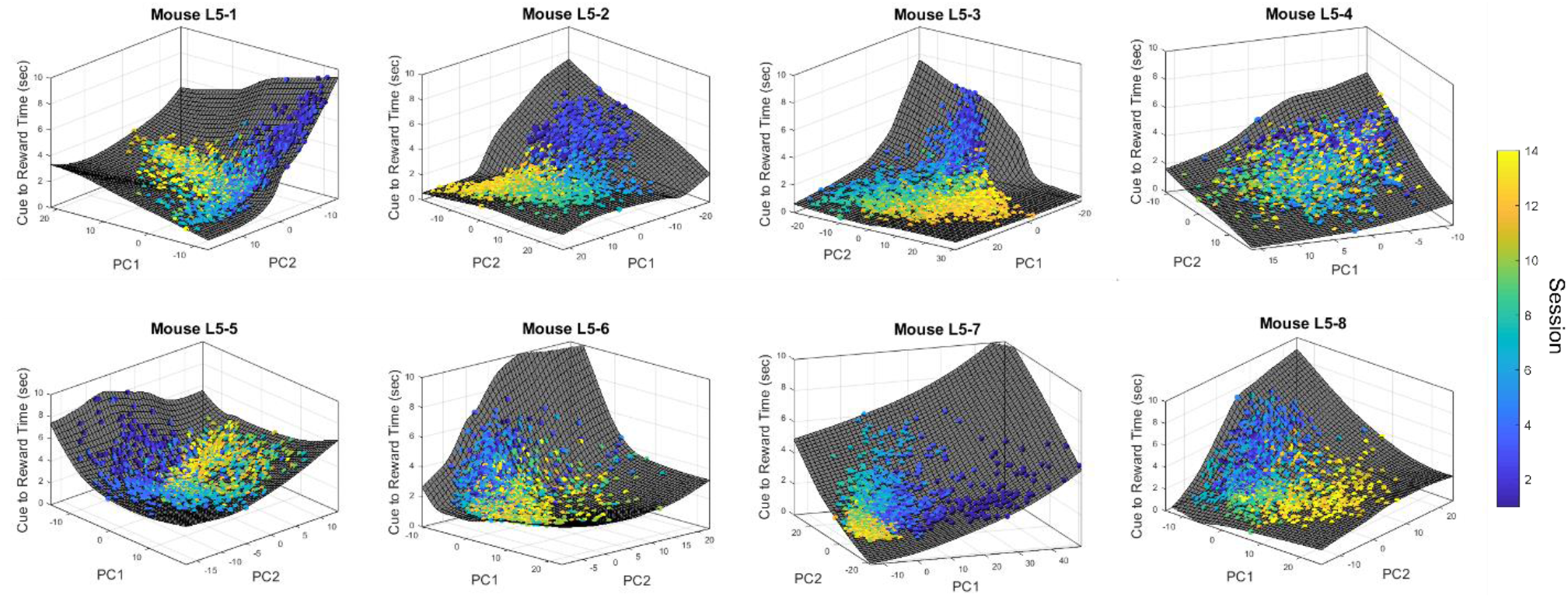
L5 connectome activity versus cue-to-reward time for rewarded trials. Each circle represents a rewarded trial and its color denotes the session number. The surface interpolation was done using MATLAB function `cftool`. The cue-to-reward time of each trial is projected onto the surface for visualization purposes.

### Connectome rewiring revolves around a core set of movement-associated neurons

In the following, we analyzed further the rewiring among neurons in L2/3 that are most significantly impacted by learning. For this purpose, we identified functional connectivity edges that have the most highly positive and negative loadings to PC1 of the connectome, as these reflect connectivity that is stably and significantly strengthened and weakened with learning, respectively (see Methods). From these edges, we obtained two groups of neurons: one group of neurons that are incident to the top positive PC1 connectivity edges and another incident to the top negative PC1 edges. Note that these groups are not mutually exclusive (*i*.*e*., there are neurons that belong to both groups). We analyzed the association of these neurons with lever movement by quantifying how much movement-related neurons—defined as neurons that have significantly higher activity during movement epochs than during non-movement epochs (see Methods)—contribute to each group. Specifically, we evaluated the fractions of neurons in each group that are movement-related in different learning phases: session 1-2 (naïve), 3-5 (early), 6-9 (mid), and 10-14 (expert). As shown in **Fig. 7a**, neurons from the top positive PC1 loading become more associated with movement with learning, while those from highly negative PC1 loadings do not exhibit a clear trend.

**Figure 7.**
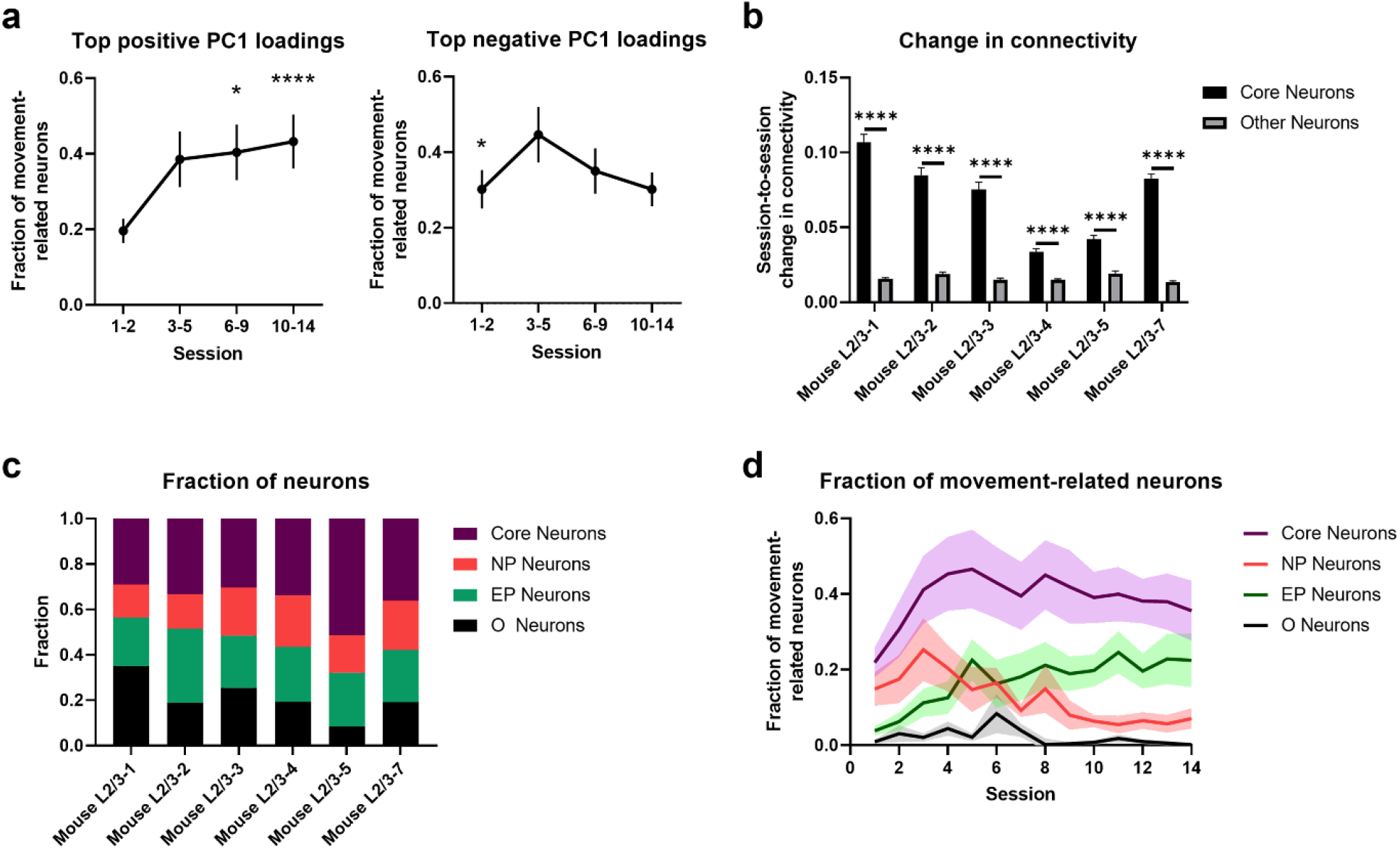
Analysis of top PC1 neurons in L2/3. **a**. Fractions of neurons from the top positive and the top negative PC1 loadings that are classified as movement related (* 5-th upper percentile, ****0.01th upper percentile). **b**. Session-to-session connectivity change among core and non-core neurons. **** *p*-value < 10^−4^ (two-sided two sample t-test). **c**. Fractions of core, NP, EP and O neurons (*n =* 6 mice). **d**. Fractions of movement-related neurons among core, NP, and EP groups. In all figures, shaded regions and error bars represent s.e.m. **(a** and **d)** across all animals or **(b)** across neurons for each animal.

In the following, we further subcategorized the two groups of neurons into three classes: Core, Naïve Phase (NP), Expert Phase (EP). Core neurons are those associated with functional connectivity belonging to both top positive and negative PC1 loadings, and as such, they represent neurons that undergo the most extensive rewiring during motor learning (see **Fig. 7b**). EP (NP) neurons are associated exclusively with the top positive (negative) PC1 loadings, and they gain (lose) functional connectivity with learning. Neurons whose edges are not associated with top PC1 loadings are called other (O) neurons. Core neurons make up 35.68 ± 3.30% (mean ± s.e.m.) of L2/3 neurons. On the other hand, NP and EP neurons comprise 18.64 ± 1.50% and 24.67 ± 1.65% (mean ± s.e.m.) of the L2/3 neuronal population, respectively (see **Fig. 7c**). Core neurons are more frequently movement related than all other groups of neurons over the learning period, as shown **Fig. 7d. Table 1** confirms that movement-related neurons are significantly overrepresented among Core neurons, but not the other classes of neurons. Meanwhile, NP (EP) neurons become less (more) movement related with learning. Therefore, the neurons that are most extensively rewired during learning are strongly associated with movement.

**Table 1.** Over-representation of movement-related neurons in Core, NP, EP, and Other neurons in L2/3. The values indicate odds ratio. \**p*-value < 0.0001 (Fisher exact test).

|  | Session 1-2 | Session 3-5 | Session 6-9 | Session 10-14 |
| --- | --- | --- | --- | --- |
| Core Neurons | 4.569* | 5.876* | 5.773* | 5.446* |
| NP Neurons | 1.494 | 0.938 | 0.531 | 0.217 |
| EP Neurons | 0.221 | 0.461 | 0.715 | 1.314 |
| O Neurons | 0.129 | 0.102 | 0.105 | 0.077 |

Looking at the connectivity more closely, Core neurons maintain a relatively stable mean degree of connectivity across the learning sessions, as shown in **Fig. 8a**, despite the fact that they are extensively rewired. This stability suggests that connectivity loss with learning is balanced by connectivity gain. Meanwhile, NP (EP) neurons show a decreasing (increasing) degree of connectivity with learning (see **Fig. 8a**). **Fig. 8b** further shows that Core neurons have a higher intragroup than intergroup connectivity, i.e., Core neurons are connected mostly among themselves. In addition, Core neurons start with a higher connectivity with NP than EP neurons, but this is reversed over the course of learning where the loss of Core-NP connectivity is countered by the gain of Core-EP connectivity. The results of the above analyses confirm the observation that learning-induced rewiring in L2/3 maintains a homeostatic number of connectivity, and further reveal that the connectome rewiring concentrates on a core group of neurons that form highly connected sub-connectome with each other.

**Figure 8.**
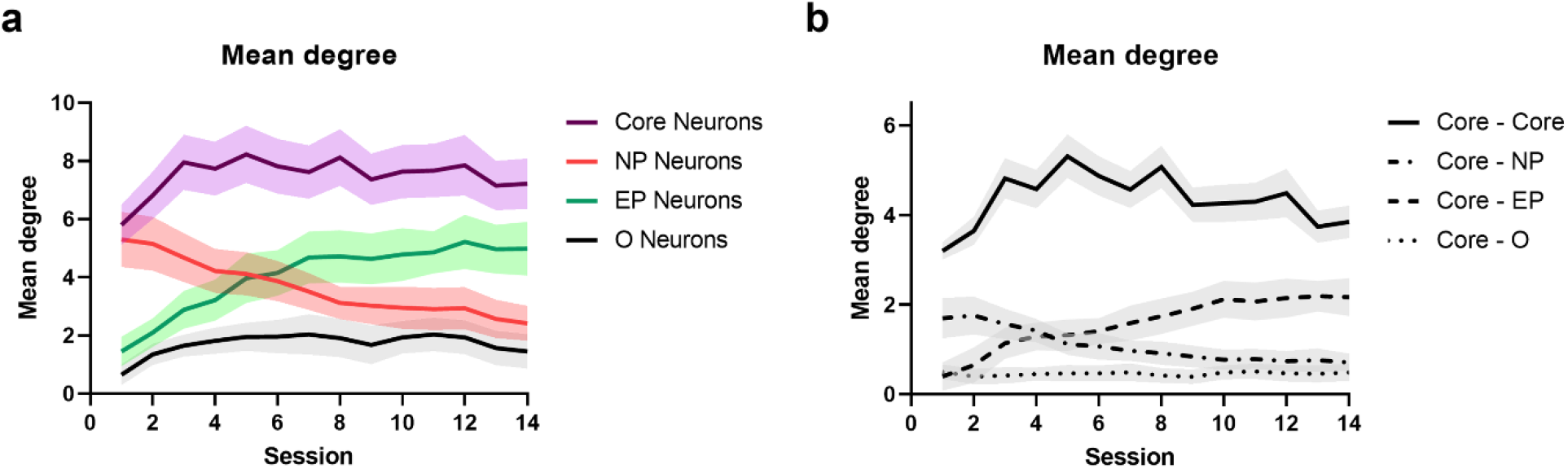
Connectivity of top PC1 neurons. **a**. Mean degree of connectivity. **b**. Mean degree connectivity of Core neurons with Core, NP, EP, and Other neurons. The shaded area denotes s.e.m. computed across all animals.

Next, we analyzed how motor skill learning affects the activity of the different neuronal groups and its relationship with movements in rewarded trials. As shown in **Fig. 9a**, Core neurons maintain a stable mean activity over the entire learning period that is relatively higher than the other groups of neurons (**Fig. 9a**). NP and EP neurons follow an opposite trend in mean activity: a decrease for NP and an increase for EP (**Fig. 9a**). The activity of Core and EP neurons become more reproducible with learning, while NP and O neurons do not (**Fig. 9b**). As described in the original study (Peters et al., 2014), learning leads to an emergence of reproducible neuronal activity in L2/3 and stereotyped movement, such that movements that are more similar to the stereotyped pattern are associated with neuronal activities that are also more similar to the learned activity pattern (Fig. 3b in Peters *et al*. (Peters et al., 2014) and **Fig. 9c**). **Fig. 9d-g** shows that such learning-associated activity-movement relationship also emerges for the above groups of neurons, but it is the most prominent for Core neurons, followed by, in decreasing degree, EP, NP, and O neurons. Interestingly, even during the initial (naïve) phase of learning, the activity of Core neurons already resembles the learned activity pattern more closely when movements are highly correlated with the stereotyped expert pattern (movement correlation > 0.5), when compared with the other groups of neurons (see **Fig. 9h**). The analysis above shows that Core neurons and their rewiring play a significant role in the emergence of stereotyped activity and movement patterns with motor skill learning.

**Figure 9.**
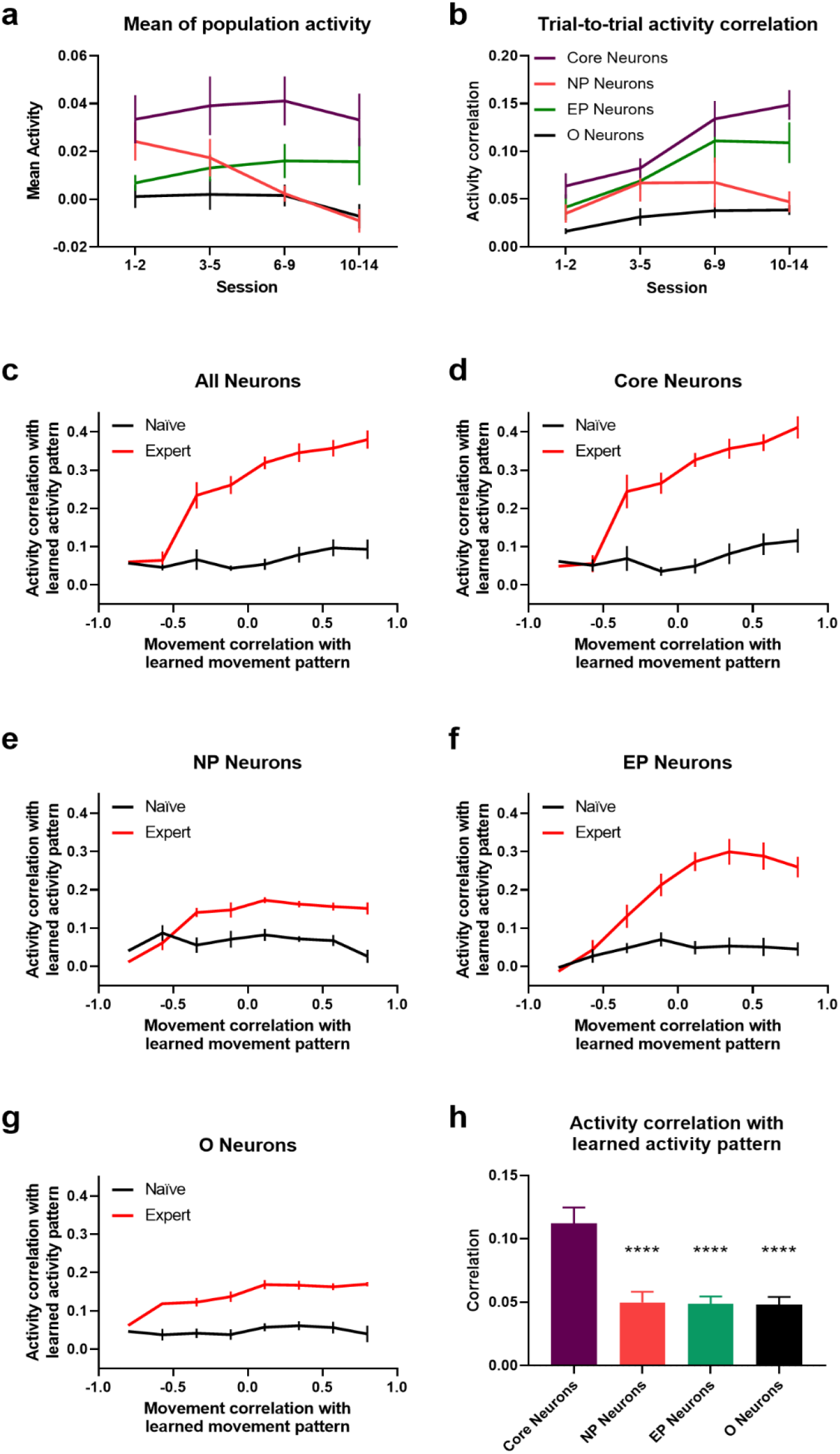
Neuronal activity and movement encoding. **a**. Mean neuronal activity of different groups in learning trials. **b**. Trial-to-trial activity correlation for different groups of neurons. (**c**-**g**) Correlations of activity vs. movement with respective expert patterns in cued trials (84-frame window) for all (**c**), Core (**d**), NP (**e**), EP neurons (**f**), and Other neurons (**g**). Naïve refers to session 1-3 and expert to session 10-14. **h**. Neuronal activity correlation with learned pattern for expert-like movements (corr. > 0.5) in naïve sessions. **** *p*-value < 0.0001 (one-sided paired t-test). Error bars indicate s.e.m computed across all L2/3 animals.

Finally, we investigated the differences across the groups of neurons in terms of encoding movement in their neuronal activity. To do so, for individual groups of neurons, we trained linear decoders that predict lever positions (lever voltages) from their neuronal activity (thresholded and smoothened spikes) for every session—called session decoders (see Methods). **Fig. 10a** shows the correlations between predicted and actual lever positions for test data that were not used in the training of the session decoders. Session decoders for Core neurons have the highest correlations, implying that Core neurons encode movements in their activity more strongly than the other groups of neurons. Besides, learning generally improves the accuracy of session decoders, indicating that there is an increase of movement-encoding within the neuronal activity of all groups of neurons in L2/3. The most significant learning-induced improvement is for EP neurons, followed by Core neurons, with only a small increase for NP and O neurons. We also tested the ability of linear decoders trained using data from the expert phase (session 10-14)—called expert decoders—to predict movements (lever positions) in the naïve phase (session 1-3). The correlations from these expert decoders are expectedly lower than using the respective session decoders (comparing **Fig. 10a** and **10b**). Still, expert decoders of Core neurons predict movements in the naïve phase significantly better than expert decoders of the other groups (*p-*value < 10^−10^), and comparable to the session decoders from each respective group. This result points to the stability of movement encoding by Core neurons over the course of learning, suggesting that learning-related rewiring might be centered around producing a stable activity pattern in a core set of neurons that maintain reliable relationship with movements.

**Figure 10.**
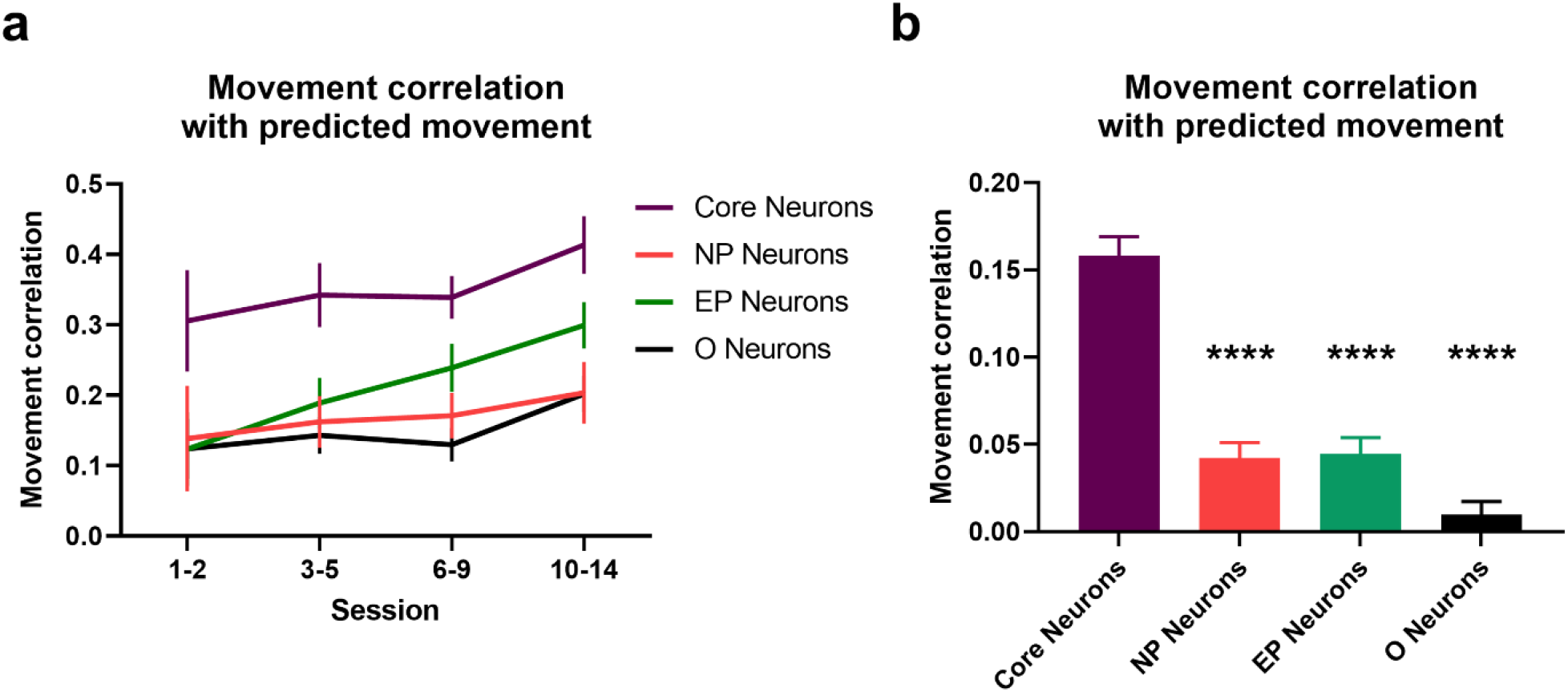
Encoding of movement in neuronal activity via linear decoders. **a**. Correlations between actual and predicted movement based on linear decoders built for each session and used to predict movements of the unseen trials of the same session. **b**. Correlations between actual and predicted movement for naïve sessions (session 1-3) using a linear decoder trained using data from expert sessions (session 10-14). **** *p*-value < 0.0001 (two-sided two-sample t-test). Error bars indicate s.e.m computed across animals.

## Discussion

Motor cortex plays a critical role in motor skill learning for integrating sensory information, establishing a motor plan, and executing movements (li and Waters, 1991; Pronichev and Lenkov, 1998; Ferezou et al., 2007; Tennant et al., 2011; Guo et al., 2015; Li et al., 2016; Makino et al., 2016; Hwang et al., 2019a, 2019b; Sauerbrei et al., 2020). In our previous studies, we demonstrated that lever-press task learning reorganizes the neuronal population activity in L2/3 and L5 of the mouse M1, as measured by *in vivo* two-photon calcium imaging. In L2/3 neurons, we observed the emergence of consistent spatiotemporal population activity accompanying learned movements (see **Fig. 2c**), leading to a stronger correlation between movement similarity and neuronal activity similarity (Peters et al., 2014). In L5, despite the same motor learning as L2/3, no consistent neuronal population activity was observed (see **Fig. 2d**), but instead dissimilar movements became further decorrelated (Peters et al., 2017). The contrasting learning-induced reorganizations of neuronal activity across different layers in the primary motor cortex suggest differences in their roles and functional objectives during motor learning (Peters et al., 2014, 2017).

Interestingly, despite the overt differences in how lever-press task learning affects the neuronal population activities in L2/3 and L5, their functional connectomes—as measured by partial correlations of Ca^2+^ spikes—show similar rewiring trajectories (see **Fig. 3 & 4**). Specifically, our analysis shows that the functional connectomes in both layers rewire to give a more interconnected structure (increased co-firing of neurons) during the first few learning sessions, and functional connections are gradually pruned in the later sessions. This result agrees well with a transient increase of L2/3 spine density measured in mice undergoing the same motor task learning, suggesting that changes of functional connectivity in L2/3 are connected, at least partly, to learning-related plasticity of dendritic spines (Peters et al., 2014). Meanwhile, the rewiring in L5 functional connectomes is associated with more subdued and delayed changes in network interconnectedness when compared to L2/3 functional connectomes: L5 peaking at around the 7^th^ learning session vs. L2/3 peaking at roughly the 3^rd^ learning session (see **Fig. 3d and 4d**). While it is known that L2/3 neurons project to L5, the lack of coherence in learning-induced changes of neuronal activity patterns and the differences in magnitude and speed of rewiring dynamics between the two layers suggest a weak inter-layer coordination of neuronal communications. That is, while the functional connectome rewiring in L2/3 and L5 occurs relatively independently of each other, both appear to engage the same rewiring principle.

Our analysis further strengthens the overt connection between spatiotemporal neuronal population activity in L2/3 and lever movement. By quantifying the functional connectome activity on a trial-to-trial basis, we demonstrated a strong correlation between functional connectome activity changes across trials with improved motor performance, as measured by the cue-to-reward time, in the early learning sessions (session 1-4, see **Fig. 5**). This period coincides with a sharp increase in the interconnectedness of the functional connectome. Thus, L2/3 functional connectomes appear to initially adopt higher interconnectivity when optimizing for motor performance. In the second half of the experiments (session 8-14), functional connectomes and their activities continue to rewire, where functional connectivity is gradually pruned, even after the cue-to-reward time reaches a plateau. On the other hand, the correlation between the changes in the functional connectome activity and motor performance is visibly less prominent in L5 when compared with L2/3 (see **Fig. 6**).

The results of these analyses give evidence for a universal learning-induced rewiring dynamics that L2/3 and L5 layers of mouse primary motor cortex engage. The overlap of time periods when L2/3 functional connectomes sharply increase their degrees of connectivity and when the cue-to-reward times markedly improve, suggests that higher interconnectedness has a functional relevance in terms of motor performance. The overt higher level of connectivity in the functional connectomes is consistent with an exploration of network state space—the space of network structures—to optimize a learning-related functional objective. Once the learning-related objective reaches an optimal (desirable) level, the functional connectome rewiring dynamics engages a different trajectory to return to homeostatic level of network connectivity and presumably a network state with high energetic efficiency. Thus, functional connectome rewiring appears to be driven by multiple objectives, where the dominant objective switches from one to another across learning and/or these objectives have large time-scale separation.

Further analyses of L2/3 neurons whose functional connectivity are most impacted by learning show how learning-induced rewiring clusters around a core set of movement-associated neurons (Core neurons). Core neurons form a highly interconnected hub in the connectome where these neurons highly interconnected among each other. Learning heavily adds and removes connectivity among these neurons, but in a manner that maintain the overall degree of connectivity, i.e., losses of connectivity are balanced by gains. Further, Core neurons reliably encode movements in response to cues even in the naïve phase, and learning-associated rewiring further improves their movement encoding. Beside the aforesaid Core neurons, there are also neurons that steadily gain and lose connectivity with learning, i.e., EP and NP neurons, respectively. While the gain of connectivity among EP neurons is associated with increased association of their activity with movement, the loss of connectivity among NP neurons does not significantly affect their activity-movement relationship. These distinct patterns of connectivity changes among the three different neuronal groups and their relationship to movements suggest that motor learning is accompanied by neuronal rewiring that reinforces the activity pattern of Core neurons that reliably encode the learned movement.

## Acknowledgments

The authors would like to acknowledge funding support from NSF-HDR IDEAS Lab (funding # 1939987, 1940202, 1940162, 1939999, and 1939992).

## Notes

**Conflict of interest:** The authors declare no competing financial interests.

### Competing Interest Statement

The authors have declared no competing interest.

